# Advancing a Science for Sustaining Health: Establishing a Model Health District in Madagascar

**DOI:** 10.1101/141549

**Authors:** Matthew H. Bonds, Andres Garchitorena, Laura Cordier, Ann C. Miller, Margaret McCarty, Benjamin Andriamihaja, Josea Ratsirarson, Andriamihaja Randrianambinina, Victor R. Rabeza, Karen Finnegan, Thomas Gillespie, Patricia A. Wright, Paul E. Farmer, Tara Loyd, Megan B. Murray, Robin M. Herrnstein, James R. Herrnstein, PIVOT Impact Team, Djordje Gikic, Mohammed A. Ouenzar, Lara Hall, Michael L. Rich

**Affiliations:** Department of Global Health and Social Medicine, Harvard Medical School, Boston, MA USA; PIVOT, Ranomafana, Madagascar/ Boston, MA USA; Department of Medicine, Stanford School of Medicine, Stanford CA USA; Ministry of Health, Madagascar; Division of Global Health Equity, Brigham and Women’s Hospital, Boston, MA USA; MICET, Antananarivo, Madagascar; Madagascar National Institute of Statistics; Department of Anthropology, Stony Brook University, Stony Brook, NY USA; Department of Environmental Studies, Emory University, Atlanta, GA USA; Renaissance Technologies, Stony Brook, NY USA; Bloomberg School of Public Health, Johns Hopkins University, Baltimore, MD USA

## Abstract

**Objective:** We demonstrate a replicable model health district for Madagascar. The governments of many low-income countries have adopted health policies that follow international standards, and yet there are four hundred million people without basic access to primary care. Closing this global health delivery gap is typically framed as an issue of scale-up, accomplished primarily through integrating international donor funds with broad-based health system strengthening (HSS) efforts. However, there is no established process by which healthcare systems measure improvements at the point of service and how those, in turn, impact population health. There is no gold standard, equivalent to randomized trials of individual-level interventions, for health systems research. Here, we present a framework for a model district in Madagascar where national policies are implemented along with additional health system interventions to allow for bottom-up adaptation.

**Setting:** The intervention takes place in a government district in Madagascar, which includes 1 district hospital, 20 primary care health centers, and a network of community health workers.

**Intervention:** The program simultaneously strengthens the WHO’s six building blocks of HSS at all levels of the health system within a government district and pioneers a data platform that includes 1) strengthening the district’s health management information systems; 2) monitoring and evaluation dashboards; and 3) a longitudinal cohort demographic and health study of over 1,500 households, with a true baseline in intervention and comparison groups.

**Conclusion:** The integrated intervention and data platform allows for the evaluation of system output indicators as well as population-level impact indicators, such as mortality rates. It thus supports field-based implementation and policy research to fill the know-do gap, while providing the foundation for a new science of sustaining health.

**Data Sharing Statement:** Data can be made available upon request by emailing.

## Introduction

The past two decades witnessed an unprecedented evolution in global health broadly connected to the Millennium Development Goals. A tripling of global health aid during the first decade of the millennium corresponded to major improvements in health outcomes for developing countries^1,2^. Yet four hundred million people continue to lack access to basic primary health care, and there remain persistent gaps in the coverage of a wide variety of essential health services. This failure is known as the global health delivery gap (or the “know-do” gap)^3,4^, and is primarily attributed to a failure in scaling up services and technologies. However, problems in scale-up are often due to breakdowns in basic implementation, which requires integrating with local health systems that are often weak^5^. As a result, large portions of the world continue to suffer from lack of basic primary care services.

Even for health system interventions that have been successful, there is an enormous challenge to scientifically describe the complex processes that led to progress and establish research systems for studying them. Though most global financing for health continues to be directed towards specific health focus areas (or “vertical” programs) such as HIV, TB, and malaria, there is a growing emphasis on the central importance of integrated health system strengthening (HSS)^6–8^. Vertical programs, which are narrowly defined, can be readily tested via controlled trials. However, there are no general standards of research on the process of strengthening health systems at the point of care, and the quantity of such research is scant^9,10^. With a few recent exceptions^11^, evidence of the effect of system interventions on population-level heath impacts, such as mortality rates, is correspondingly negligible and controversial^12–14^. As the Sustainable Development Goals articulate the world’s more expansive aspirations for human development, there is ever greater need for quality evidence on how best to improve the systems that sustain human life^15^. Here, we present a framework for a model district health system in Madagascar.

Madagascar is one of the poorest countries in the world^16^. After a coup in 2009, the country became ineligible to receive international aid during a period of unprecedented international investment in global health. Official Development Assistance to Madagascar dropped from a high of 71 USD per capita in 2004 to 17 USD per capita in 2012, among the lowest in Africa^17^. In 2014, total per capita spending on healthcare in Madagascar was the lowest in the world, at14 USD ^17^. Since January 2014, due to democratic elections, Madagascar’s newly recognized government has become eligible for official aid from foreign governments, creating a singular opportunity for transforming the national health system.

Our initiative in Madagascar is based on a partnership between the Madagascar government (i.e., Ministry of Health) and the nongovernmental organization PIVOT, and is being carried out in Ifanadiana District, located in the southeast of the country. This initiative is modeled in part on a recent HSS initiative in Rwanda^18^, which oversaw some of the fastest known reductions in under-five mortality in the world^11,19^. The goal is a rapid expansion of health services at all levels of the Ministry of Health (MoH) system within a government district (community, health center, and hospital). This is done across the WHO’s six building blocks of HSS: service delivery, health workforce, information systems, medicines and supplies, financing, and leadership/governance. The health interventions are guided by MoH policies and include a new platform for information systems and research that includes: 1) government health management information systems (HMIS); 2) real-time monitoring and evaluation; and 3) a geocoded longitudinal cohort demographic and health study of 1,520 households (~9,000 individuals) with baseline established prior to the start of the intervention. The data platform thus allows for the evaluation of system output indicators and population-level impacts such as mortality rates, critical for generating transferable (and externally validated) lessons for HSS in a quasi-experimental context.

## District-level Health System Strengthening Intervention

Systemic failures in any one building block of HSS or at any of the levels of the health system have cascading effects that weaken the system overall. Although “health systems” are often conceptualized as national, a government district is an important unit of intervention where the majority of patients experience the continuum of health care services^21^. It is a common unit for piloting health initiatives intended for national scale-up. In Madagascar, as in most developing countries, there is one hospital per district (~200,000 people), each with 30–50 beds (Madagascar has 114 districts total). Most patients are referred to the district hospital from basic health centers, which are primary care facilities that serve the population of each commune (~10,000 people, ~10–20 per district). In addition to providing primary health care, basic health centers supervise community health workers (CHWs), who are locally elected volunteers. In Madagascar there are 2 CHWs per village area, which is called a fokontany (~10 per commune) – the smallest administrative unit of the government. CHWs are responsible for diagnosis and treatment of the most common causes of under-five mortality (i.e., diarrhea, pneumonia, and malaria) and serve to link pregnant mothers with prenatal care.

**Figure 1.**
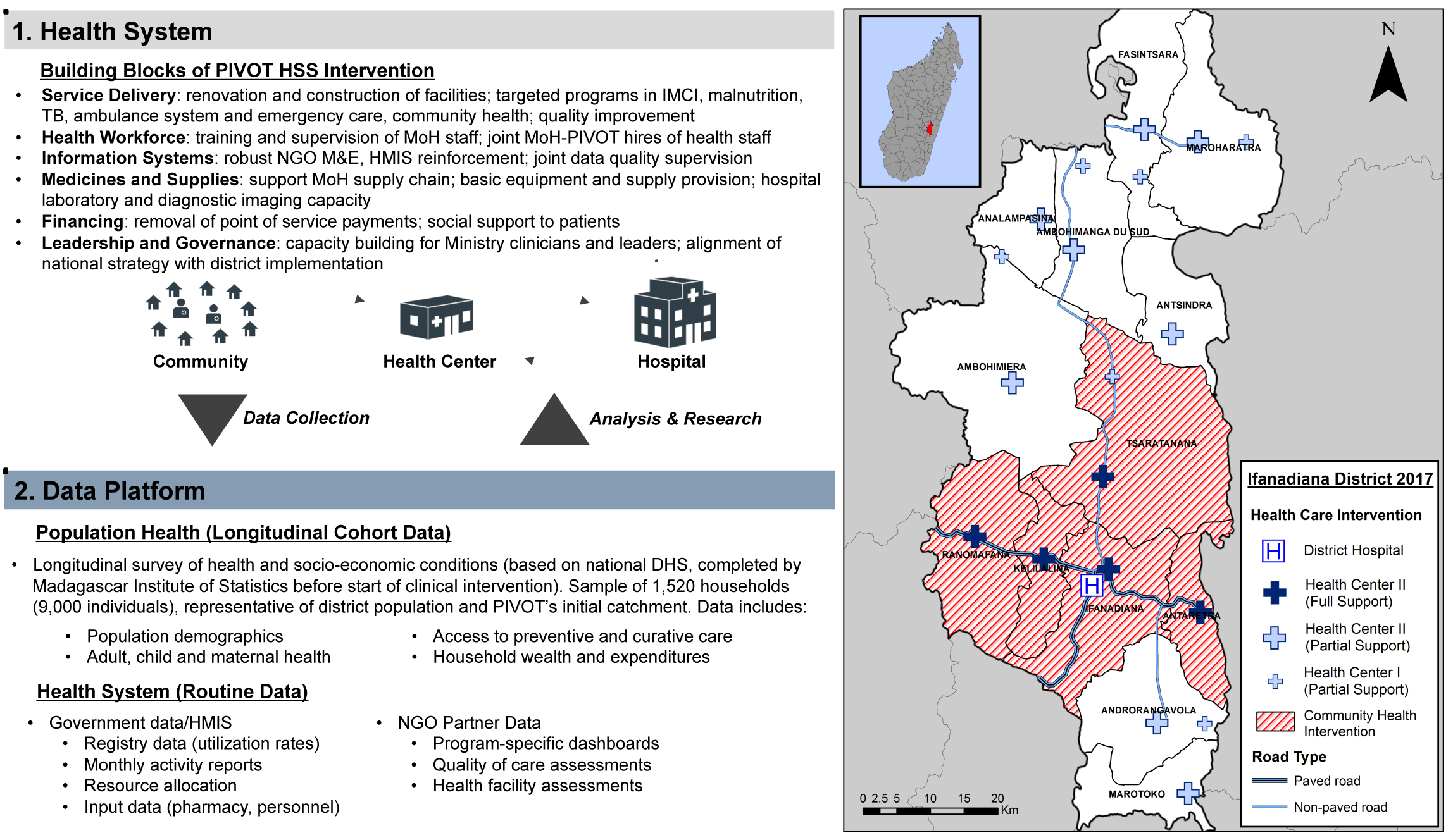
A New District Platform for Evidence-Based Health System Strengthening. The platform, a partnership between the Ministry of Health, nongovernmental and academic partners, provides a feedback system between 1) health system interventions at the community, health center and hospital levels across a government district, and 2) information systems of routine data and a population-based longitudinal cohort study^20^. The right panel is a map of the interventions in Ifanadiana District; there is a Health Center II per commune, each with a medical doctor. Health Center I does not having a medical doctor.

PIVOT-MoH interventions at the hospital, health center, and community levels include infrastructure upgrades, expanding quantity and capacity of personnel, improving supply chain management, providing medical equipment, reducing user fees, malnutrition treatment, social programs, information systems, and ambulance services, among others. Figures 2 and 3 provide two examples (malnutrition treatment and medicine financing) of how specific programmatic interventions are integrated with the data platform to inform the process of implementing and scaling the intervention.

## Data Platform

There is a dearth of evidence on how health interventions change population-level indicators. For randomized trials of specific interventions, before and after data are routinely collected, but the interventions do not typically cause measureable population-level impacts. Health system strengthening efforts are often thought to have population-level impacts, but their scale usually precludes a counter-factual: there is rarely true baseline data for both intervention and comparison groups. Our platform aims to fill this gap^20,22^.

The data platform provides the foundation for understanding underlying health conditions and allows for measurement of the effect of interventions on health system processes (or outputs) and on human health outcomes. There are three main components.

1. The national health management information system (HMIS) captures aggregate registry data on community, health facility and hospital activities, including patient visits per condition, health care utilization, pharmacy management, logistics and capacity at all levels. Due to improved data quality, which is often considered unreliable, PIVOT and the MoH jointly conduct supervised data quality assessments, as well as IT and database training.
2. Monitoring and evaluation data are featured on dashboards for PIVOT programs including real-time analyses of costs per patient, stock-out rates, and patients lost to follow-up, among other input, output and outcome indicators. These data are collected from PIVOT project records and facility-based data sources.
3. In collaboration with the Madagascar National Institute of Statistics, a district-wide longitudinal cohort study was initiated with a representative sample of households drawn in two strata—PIVOT’s initial catchment area (population 80,000) and the rest of the district (population 100,000). The study is based on the Demographic and Health Survey (DHS), which captures data on health, health care access, mortality, and socioeconomic status among other information. Baseline status of 1520 households was collected in 2014 before the initiation of the intervention activities. A second round of data collection occurred in 2016, and further rounds will be repeated every two years. Because the same survey (the DHS) is conducted routinely at the national level (roughly every five years), it is possible to compare populations within the district as well as across the rest of Madagascar. The protocol for the longitudinal cohort study is available in the Supporting Information.

**Figure 2.**
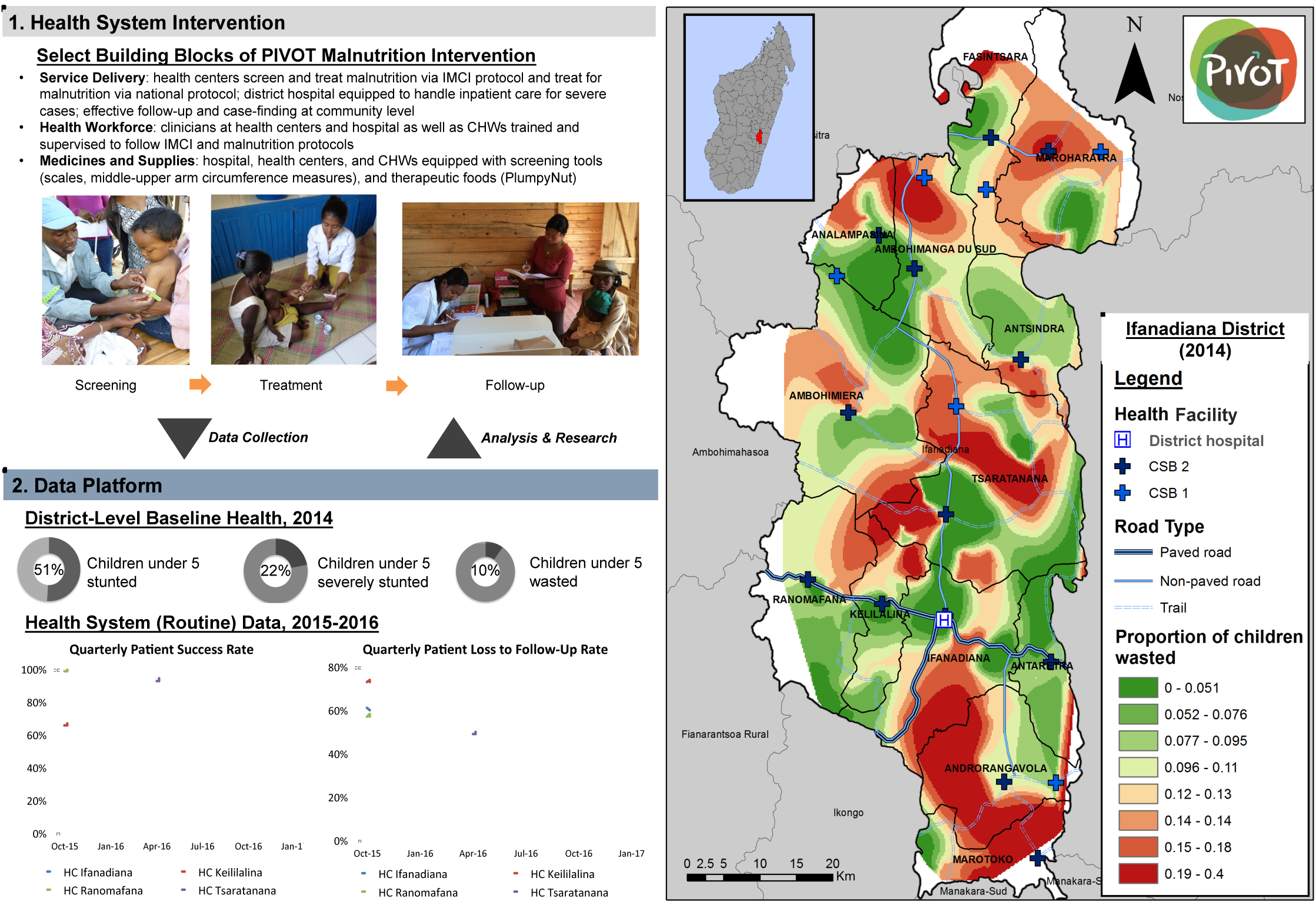
Malnutrition Program. The process of strengthening health systems occurs through a series of individual initiatives, such as 1) a malnutrition program, which PIVOT-MoH implements at the community, health center, and hospital levels in the government district. Implementing malnutrition protocols requires simultaneously implementing general child health protocols (i.e., integrated Management of Child Illness (IMCI). Thus, implementing one child health initiative aligned with national policies can serve to strengthen broader systems for child health. 2) Data are routinely collected to analyze the effectiveness of the malnutrition program. 3) The longitudinal cohort study indicates spatial distribution of malnutrition in the district. These data are monitored over time to inform program priorities for measuring impacts.

In addition to these databases, a fundamental source of information is contextual knowledge. Improving health systems requires identifying problems as they arise and adapting to them. The causes of policy failures depend on local context, and can range from resource constraints, to technical capacity, corruption, technical efficiencies, or bottlenecks in financing. Information on such barriers is often absent from routine data collection systems, but obvious to local practitioners, community members and patients. In addition to performing targeted qualitative studies, being directly involved in field-based service provides critical opportunities for identifying such breakdowns in the system and addressing them.

**Figure 3.**
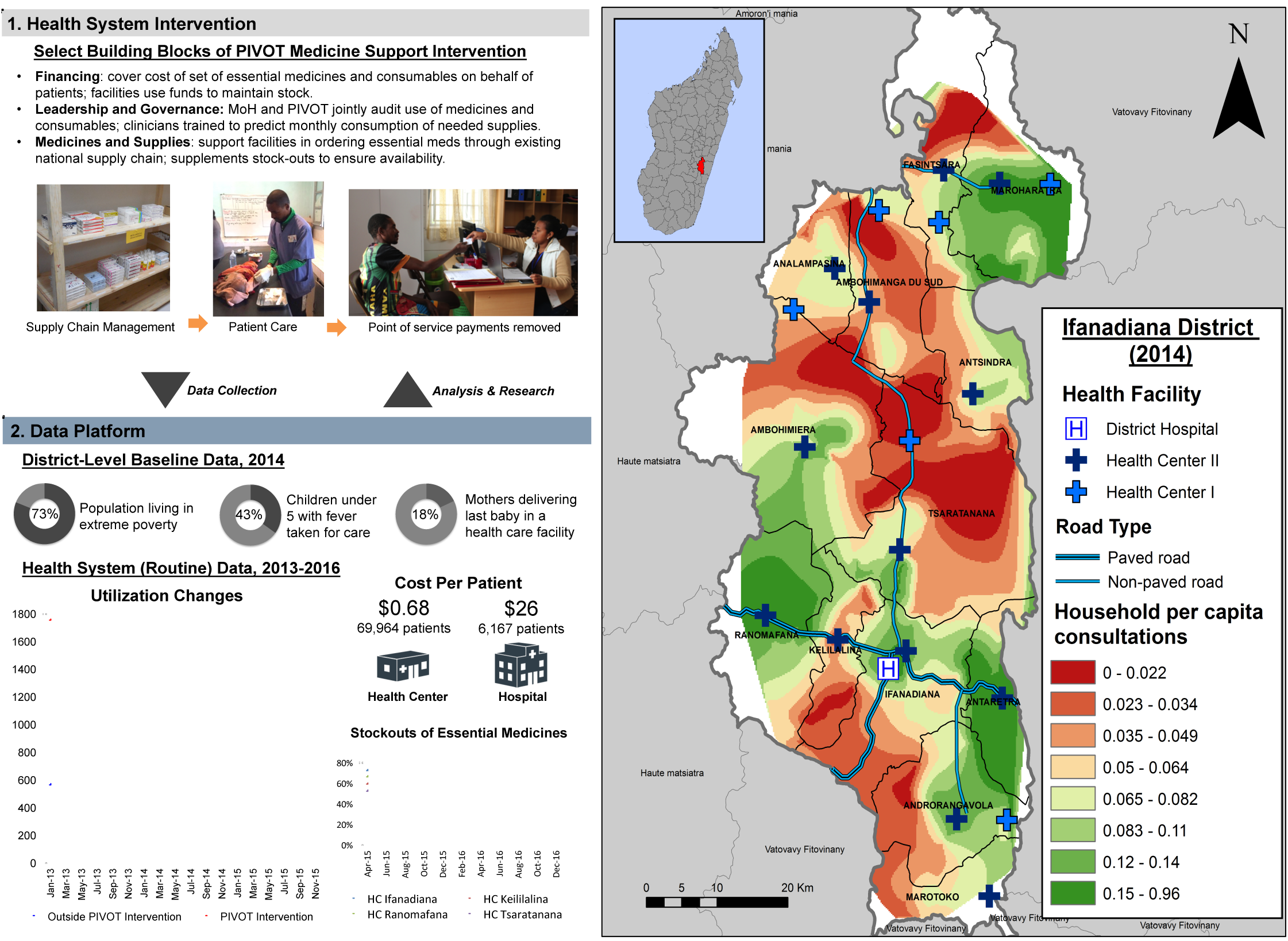
Medicine Support Program. 1) Through a system of reimbursing health facilities for patient costs of 40 medicines and 20 consumables, point-of-service payments are effectively removed. 2) This has resulted in immediate increases of health center utilization at an incremental cost of around $.68 per patient ($26 per patient in the hospital). The right panel presents of the spatial distribution of changes to reported consultations based on household surveys from 2014–2016^22^.

## Discussion

We suggest that he persistence of the global health delivery gap is attributable in part to the distance between 1) health care providers and implementers who specialize in providing care for individuals, and 2) policymakers and analysts engaged in aligning international funding with national health strategies. Although significant progress has been made, there is a general trade-off between the complexity and scale of interventions and the ability to create data systems that allow for inferring impacts over time.

Over the past decade a number of initiatives across Africa have taken root that attempt to fill this gap through various HSS strategies implemented at a population scale. Many of these initiatives - notably in Kenya, Ghana, Mozambique, Rwanda, Tanzania, Malawi, Mali, and Zambia - are integrated with data systems^23–25^. However, to our knowledge, there are yet to be statistically rigorous findings that under-five mortality rates have improved as a result of the health system intervention, though results in Rwanda are highly suggestive. Thus there still remain basic questions on how health systems can be improved at the point of service to create population level change.

Our work builds on these other initiatives by creating interventions across all of the building blocks of HSS at the levels of the health system at a scale that is large enough to fully alter health conditions for the population, but small enough to measure and generate transferable knowledge for scale. This has the potential to become the first initiative to reveal statistically rigorous differences (i.e., difference-in-differences) in under-five mortality rates as a result of the intervention. In addition to supporting modern health care delivery, such an integrated service and data platform can provide a foundation for basic scientific inquiry on causes of disease as well as for innovation that together will contribute to the evolving science of sustaining health.

## Authors’ Contribution Statement

Conception and design of the study was done by MHB, AG, LC, ACM, MGM, PEF, MM, PCW, BA, TL, JR, RMH, JRH, DG, MAO, LH, MLR, Data collection was done by MHB, AR, AG, LC, ACM, VRR, KF, PIT, DG, LH, All authors contributed to writing of the manuscript. All authors have approved the version of the manuscript for publication.

## Acknowledgments

Core concepts of this initiative were generated from a PHIT grant from the Doris Duke Charitable Foundation to MLR. M.H.B. was funded by a James McDonnell Foundation grant (no. 220020322) and NIH grant (no. K01TW008773) from the Fogarty International Center and from a grant from the Herrnstein Family Foundation. We are grateful for helpful feedback from Nancy Ferguson, Robert Cunningham, and Steve Luby. The PIVOT Impact Team includes: Herinjaka Andriambolamanana, Hassan Bouziane, Elliane S Hery, Lillie Malonzeu, Charles Meade, Luc Rakotonirina, Ranjoto, Faramalala Rabemananjara, Ando Randrianandrasana, Tahiry Raveloson, and Anicet Tsifahera. Special thanks to PIVOT staff, Madagascar Ministry of Health and Centre ValBio at Stony Brook University.

## Data Sharing Statement

Data can be made available by contacting.

